# Co-regulation of hormone receptors, neuropeptides, and steroidogenic enzymes across the vertebrate social behavior network

**DOI:** 10.1101/435024

**Authors:** Brent M. Horton, T. Brandt Ryder, Ignacio T. Moore, Christopher N. Balakrishnan

## Abstract

The vertebrate basal forebrain and midbrain contain a set of interconnected nuclei that control social behavior. Conserved anatomical structures and functions of these nuclei have now been documented among fish, amphibians, reptiles, birds and mammals, and these brain regions have come to be known as the vertebrate social behavior network (SBN). While it is known that nuclei (nodes) of the SBN are rich in steroid and neuropeptide activity linked to behavior, simultaneous variation in the expression of neuroendocrine genes among several SBN nuclei has not yet been described in detail. In this study, we use RNA-seq to profile gene expression across seven brain regions representing five nodes of the vertebrate SBN in a passerine bird, the wire-tailed manakin *Pipra filicauda*. Using weighted gene co-expression network analysis (WGCNA), we reconstructed sets of coregulated genes, revealing striking patterns of variation in neuroendocrine gene expression across the SBN. We describe regional expression variation networks comprising a broad set of hormone receptors, neuropeptides, steroidogenic enzymes, catecholamines, and other neuroendocrine signaling molecules. Our findings highlight how heterogeneity of brain gene expression across the SBN can provide functional insights into the neuroendocrine and genetic mechanisms that underlie vertebrate social behavior.

## INTRODUCTION

Over the last two decades, our understanding of the neuroendocrine and genetic bases of vertebrate social behavior has progressed rapidly due to integrated studies of how hormones and gene expression interact to modulate brain function and, ultimately, behavior.^1,2^ Much of this research has focused on well-defined neural circuits that are highly conserved across vertebrate taxa, such as the social behavior network (SBN). The vertebrate SBN comprises six reciprocally connected brain regions, or nodes, located in the basal (i.e., ‘limbic’) forebrain and midbrain.^3,4^ These nodes are: 1) the extended medial amygdala, including the bed nucleus of the stria terminalis (BSTm) and, in birds, nucleus taenia (TnA); 2) the lateral septum (LS); 3) the preoptic area (POA); 4) the anterior hypothalamus (AH); 5) the ventromedial hypothalamus (VMH); and 6) the midbrain central grey (CG), including the intercollicular nucleus (ICo). Mounting evidence from multiple vertebrate taxa suggests that these nodes are activated to varying degrees during social interactions, and that collective neural activity across these nodes plays an essential role in regulating various forms of social behavior (e.g., sexual behavior, aggression, parental care, and social affiliation).^3–7^ Because the SBN represents a fundamental element of the neural machinery underlying vertebrate social behavior, the brain regions that make up this network are prime focal points for investigating how changes in neuroendocrine activity and gene expression modulate intraspecific and interspecific variation in behavior.^8–10^

The sex steroids and nonapeptide hormones (e.g., vasopressin, oxytocin) are potent regulators of vertebrate social behavior. Accordingly, the brain regions of the SBN are generally rich in sex steroid and nonapeptide receptors and thus responsive to these hormones.^3,4,6,11–14^ The sex steroids and nonapeptides can have multiple receptors^15–18^ and the distribution of these receptors determines the sensitivity of the SBN nodes to hormone action. In addition, regions of the SBN are, to varying degrees, sites of steroid and nonapeptide hormone synthesis and metabolism. ^14,19^ Our understanding of how these hormones act to modulate social behavior requires a holistic approach that can quantify the patterns of neuroendocrine gene expression that dictate hormone action in the SBN, including genes for hormone receptors, steroidogenic enzymes, and nonapeptides. Moreover, steroids can act through both genomic and non-genomic pathways to affect gene expression and subsequent behavior.^15,16,20^ As such, identifying the molecular products of steroid actions that influence neuron activity in the SBN can shed much needed light on the mechanistic underpinnings of steroid-mediated behavior.

The accessibility of RNA sequencing (RNA-seq) technology to researchers studying non-traditional model organisms provides new and exciting opportunities to deepen our understanding of the molecular pathways through which hormones influence the brain and behavior.^9,21–23^ Functional genomics provides an ideal tool for beginning to understand how suites or networks of co-expressed genes modulate behavior. To date, a number of studies have leveraged whole-brain transcriptomic data to identify the gene modules (i.e., sets of co-regulated genes) underlying among-individual variation due to sex, social status, ecotype, and reproductive strategy^22,24–26^ as well as interspecific differences in reproductive and social behaviors across species.^27–29^ Mounting evidence suggests, however, that patterns of gene co-expression vary among subdivisions of the vertebrate brain.^30–33^ For example, studies of birds and mammals have revealed distinctive gene co-expression profiles in the amygdala and hypothalamus^34–36^, suggesting that suites of behaviorally relevant genes are differentially expressed among nodes of the SBN. Indeed, results from candidate gene studies have demonstrated that expression levels of key neuroendocrine genes can vary widely across regions of the SBN, and that relationships between neuroendocrine gene expression and behavior are region-dependent.^14,36–38^ Moving forward, it is therefore necessary to examine individual nuclei (or cell groups) as opposed to whole brains to learn how patterns of gene co-expression across nodes of the SBN and other behavioral circuits underlie variation in social behavior.^9^ To our knowledge, no study has yet employed whole-transcriptome sequencing to compare gene co-expression profiles across several nuclei of the avian SBN.

To continue probing the neuroendocrine and genomic architecture of complex social behaviors, we used RNA-seq and weighted gene co-expression network analyses (WGCNA) to describe region-specific patterns of gene expression across multiple nodes of the SBN in the male wire-tailed manakin (*Pipra filicauda*). This South American lek-breeding bird exhibits a rare form of cooperation, whereby males form coalitions to perform elaborate cooperative courtship displays. These males establish long-term display partnerships that form the basis of complex social networks, and which have clear fitness benefits for both territorial (i.e., increased reproductive success)^39^ and non-territorial males (i.e., increased probability of territory acquisition).^40^ Territorial and non-territorial males show substantial variation in behavioral phenotype, yet the hormonal and genetic bases for these differences remain largely unknown. Here, we sequenced total RNA extracted from six distinct brain nuclei (BSTm, TnA, LS, POM, VMH, and ICo) representing five of the six nodes of the vertebrate SBN, and an additional brain region (arcopallium intermedium, AI) hypothesized to be involved in androgen-dependent display behavior in manakins^41^ from male wire-tailed manakins. We compared gene expression profiles among these regions to define patterns of uniqueness and congruity in gene co-expression across the SBN. In addition, we use WGCNA to describe gene networks and identify hub genes. Finally, we considered region-specific expression levels of candidate neuroendocrine genes involved in steroid and neuropeptide signaling to build a foundation for future studies of how hormone-signaling pathways modulate variation in male behavioral phenotype in this and other cooperative species. Because circulating testosterone levels vary according to male social status in wire-tailed manakins^42^, we pay special attention to those the genes involved in testosterone signaling (e.g., sex steroid receptors and steroidogenic enzymes) as well as those involved in the steroid-sensitive neuropeptide systems (e.g., vasotocin, vasoactive intestinal peptide, and their receptors).

## METHODS

### Subjects and Field Sampling

We collected tissues from four male wire-tailed manakins (*Pipra filicauda*) near the Tiputini Biodiversity Station (0°38’ S, 76°08’ W) in the Orellana province of eastern Ecuador during late December 2015, a period that coincides with peak breeding activity in this population. Males in this species can be categorized into four social classes according to plumage (age) and social status (territorial versus floater), as described by Ryder et al‥ ^39,42^ The males we sampled represented two of these classes; two individuals were definitive plumage (i.e., ≥ third year) males that held display territories on leks, and two individuals were pre-definitive plumage (i.e., second year) males known as ‘floaters’ because they did not hold display territories. Due to small sample size we did not formally compare gene expression in males based on social status.

The birds were captured in mist nets on leks between 07:30-10:00, and blood sampled before being transported to a nearby workstation where they were euthanized via decapitation. Whole brains were then extracted from the skull and rapidly frozen in powdered dry ice 4-6 minutes after euthanasia. Brains were kept frozen on dry ice for 1-4 hours in the field until they were transferred to a liquid nitrogen dry shipper (ThermoScientific) where they were stored at cryogenic temperatures for 14-16 days until imported into the United States (US). Once in the US, brains were stored at −80°C for 6.5 months until cryosectioning, microdissection, and RNA extraction (described below).

### Cryosectioning, Microdissection, and RNA Extraction

Brains were cryosectioned at −12°C to −14°C in the coronal plane consistent with the orientation used by Stokes et al. (1974). Frozen sections 150μm thick were briefly (<10 sec) thaw-mounted onto Fisherbrand Premium Superfrost microscope slides, after which they were held on dry ice until microdissection. During microdissection, slides with adhered sections were placed on slabs (15 × 15 × 3cm) of polished granite, which were kept frozen on dry ice throughout the procedure. With the aid of a stereomicroscope and fiber optic light source, seven brain regions of interest (ROIs) were located using readily identifiable landmarks (e.g., fiber tracts, ventricles) and sampled by using a frozen, stainless steel tissue corer (Fine Science Tools) to collect a 0.35, 0.50, or 0.80mm diameter punch from the ROI. The punch diameter depended on the size of the ROI (see below).

We sampled the arcopallium intermedium (AI), as delineated by Fusani et al.^41^, by taking 0.8mm punches from the left and right hemispheres of two consecutive sections. Nucleus taenia (TnA), as defined by Stokes et al.^43^, and the medial bed nucleus of the stria terminalis (BSTm), as described by Kelly et al.^44^, were sampled by taking 0.5mm punches from the left and right hemispheres of four consecutive sections. The portion of the intercollicular nucleus (ICo) medial to the nucleus mesencephalicus lateralis, pars dorsalis (MLd)^43,45^, was sampled by taking 0.5mm punches from the left and right hemispheres from four consecutive sections. We sampled the lateral septum, as delineated by Goodson et al.^46^, by taking 0.35mm sections from the left and right hemispheres in four consecutive sections. Because the ventromedial hypothalamus (VMH)^47^ and medial preoptic area (POM)^48^ are located at the midline of the brain, they were sampled by taking 0.8mm punches centered on the midline to simultaneously capture the left and right portions of these ROIs; each ROI was sampled in three to four consecutive sections. Although we also sampled the anterior hypothalamus (AH) and central mesencephalic grey (GCt), RNA yields from some samples of these regions were insufficient for sequencing and are not described hereafter.

Frozen tissue punches were placed directly into room temperature QIAzol lysis reagent (QIAGEN) and disrupted while thawing using a pellet pestle mixer and hand-held motor (Fisher Scientific). This lysate was then frozen at −80°C until the cryosectioning and microdissection of all brains were completed (≤ 3 days). The frozen tissue lysate was thawed and immediately passed through a QIAshredder spin column (QIAGEN) to further homogenize tissue lysates. We then proceeded immediately with RNA extraction and purification, which was performed using miRNeasy Micro Kits (QIAGEN) according to the manufacturer’s recommended protocol for tissue samples containing < 1μg of RNA. We included an on-column DNase I digestion step using an RNase-Free DNase Set (QIAGEN) according to the manufacturer’s protocol for tissue samples containing < 1μg of RNA. Purified RNA was eluted with 14μL of RNase-free water and stored at −80°C until quantitation and RNA integrity analysis. RNA extraction and purification for all samples was completed in a single run. Aliquots (1μL) of purified RNA samples were quantitated using a Qubit fluorometer and a Qubit™ RNA HS Assay Kit (Invitrogen). Total RNA concentrations ranged from 7.3 − 440.0 μg/mL (mean = 15.4 μg/mL). We assessed RNA integrity using the Agilent 2100 Bioanalyzer System and RNA 6000 Pico Kits (Agilent Technologies). RNA integrity numbers (RINs) ranged 7.8-9.8 (mean = 8.95); RIN values were ≥ 8.0 for all but one sample.

### RNA Sequencing & Analysis

Library preparation and sequencing were done at the University of Illinois Roy J Carver Biotechnology Center. Thirty RNA-seq libraries were prepared using the Illumina TruSeq Stranded mRNAseq Sample Prep Kit. Libraries were quantitated by qPCR and then sequenced on two lanes of an Illumina HiSeq 4000. Fastq files were generated and demultiplexed with bcl2fastq (v2.17.1.14). Adapter & quality trimming was done using TrimGalore!^49^ under default settings. We then mapped reads to the *Manacus vitellinus* reference genome (ASM171598v1) using HiSat^50^. Manakins of the *Manacus* and *Pipra* genera are relatively closely related within the Family Pipridae^51^, so we expected interspecific read mapping to work well between these two taxa. To quantify the abundance of transcripts we counted the number of reads that mapped to NCBI-predicted genes for the *M. vitellinus* genome using HT-seq.^52^ We normalized count data for analysis using the variance-stabilized transformation in DE-Seq2.^53,54^ After normalizing read counts using DE-Seq2, we used the plotPCA function in DE-Seq2 and PCAexplorer^55^ to visualize and describe overall expression profiles. PCA analysis used the top 500 genes in the dataset based on variance.

### Weighted Gene Co-expression Network Analysis & Differential Expression Testing

We sought to identify networks of coregulated genes that characterized each of the sampled nodes of the SBN. To this end, we used weighted-gene co-expression network analysis.^56,57^ Unlike PCA, WGCNA does not impose orthogonality onto gene sets, and therefore is more appropriate for describing possible regulatory pathways. Regulatory modules in WGCNA are created without *a priori* information on the sampling design. Using the variance stabilized count data from DE-Seq2, we generated a signed network by setting soft threshold power (β) = 18, minimum module size = 30, and module dissimilarity threshold = 0.30. Soft thresholding was chosen by plotting β against Mean Connectivity and selecting the point at which we observed a plateau in Mean Connectivity representing a scale-free topology. Following module reconstruction, we tested for correlations between modules and each of the seven sampled brain regions treating each as a binary variable. Because four individual birds were sampled, we also included the bird ID as a variable in the model.

We functionally characterized co-expression networks using Gene Ontology (GO) analysis using the rank order based approach in Gorilla.^58^ For each module, genes were ranked based on their module membership score determined in the WGCNA analysis. We preferred this rank-order based approach (as opposed to strict module assignment) as it reflects the correlation among modules, and because some genes could be assigned to multiple modules. GOrilla tests for enrichment towards the top of a gene list, where the “top” is defined based on the size of a particular GO category rather than an alpha value threshold. GOrilla also calculates an enrichment statistic based on N, the number of genes in the data set, B the number of genes in the whole dataset of a particular GO category, n, the number of genes in the top of the gene list and b, the number of genes that represent that GO category in the top of the gene list. Gene IDs were assigned based on annotation by NCBI and held in the gff file associated with the *Manacus* genome assembly. We then visualized the network structure for the top 500 strongest connections using a topological overlap measure and plotted the data in program R^59^ using the iGraph^60^ and ggnetwork packages. We identified module hubs as genes with more than 20 connections within this set.

All RNA sequencing data have been deposited to GenBank (PRJNA437157). The raw count matrix and code for the WGCNA analysis are available at: https://github.com/chrisbalakrishnan/PipraFilicaudaRNAseq

## RESULTS

RNA-sequencing yielded an average of 26 million reads per sample. Seventy-two percent of the trimmed reads mapped to the genome and 49% of the reads could be unambiguously assigned to predicted transcripts using htseq-count. We detected expression (1 read in at least 1 individual) for 16,796 genes, or 89.5% of the 18,775 annotated genes in the genome. As required by WGCNA, we filtered low expression genes keeping 12,765 that passed WGCNA quality control criteria. Of these, 9,598 could be linked to functional annotations in the Gene Ontology database. Only these 9,598 genes were used for GO analyses.

Principal components 1 and 2 explained 42.4% and 19.1% of the variation in the expression data, respectively (Fig. 1a). Sample expression profiles clustered by the nucleus from which they were extracted (Fig. 1a), reflecting consistency in dissection as well as regional differences in gene expression. Gene expression profiles from the two hypothalamic nuclei, POM and VMH, were associated with positive PC1 values, whereas those from the amygdala (TnA) and the adjacent arcopallium intermedium (AI) were associated with negative PC1 values. PC2 was also associated with brain region with high values for LS, intermediate values for POM, VMH, BSTm, AI and TnA, and low values for the midbrain nucleus ICo. Loadings for the genes most strongly associated with PC1 and 2 are presented in (Fig. 1b). Many of the genes with strong, positive PC1 loadings are involved in neuropeptide signaling. These include Galanin and GMAP Prepropeptide (GAL), Aldehyde Dehydrogenase 1 Family Member A1 (ALDH1A1), Neuropeptide VF Precursor (NPVF), Prodynorphin (PDYN), and Melanocortin Receptor 3R (MC3R). In contrast, genes with positive loadings on PC2 are involved with more developmental functions, including homeobox genes Distal-Less Homeobox (DLX1), DLX5, DLX6 and ISL Lim Homeobox 1 (ISL1). Neuropeptide Y (NPY) was also positively associated with PC2.

**Figure 1.**
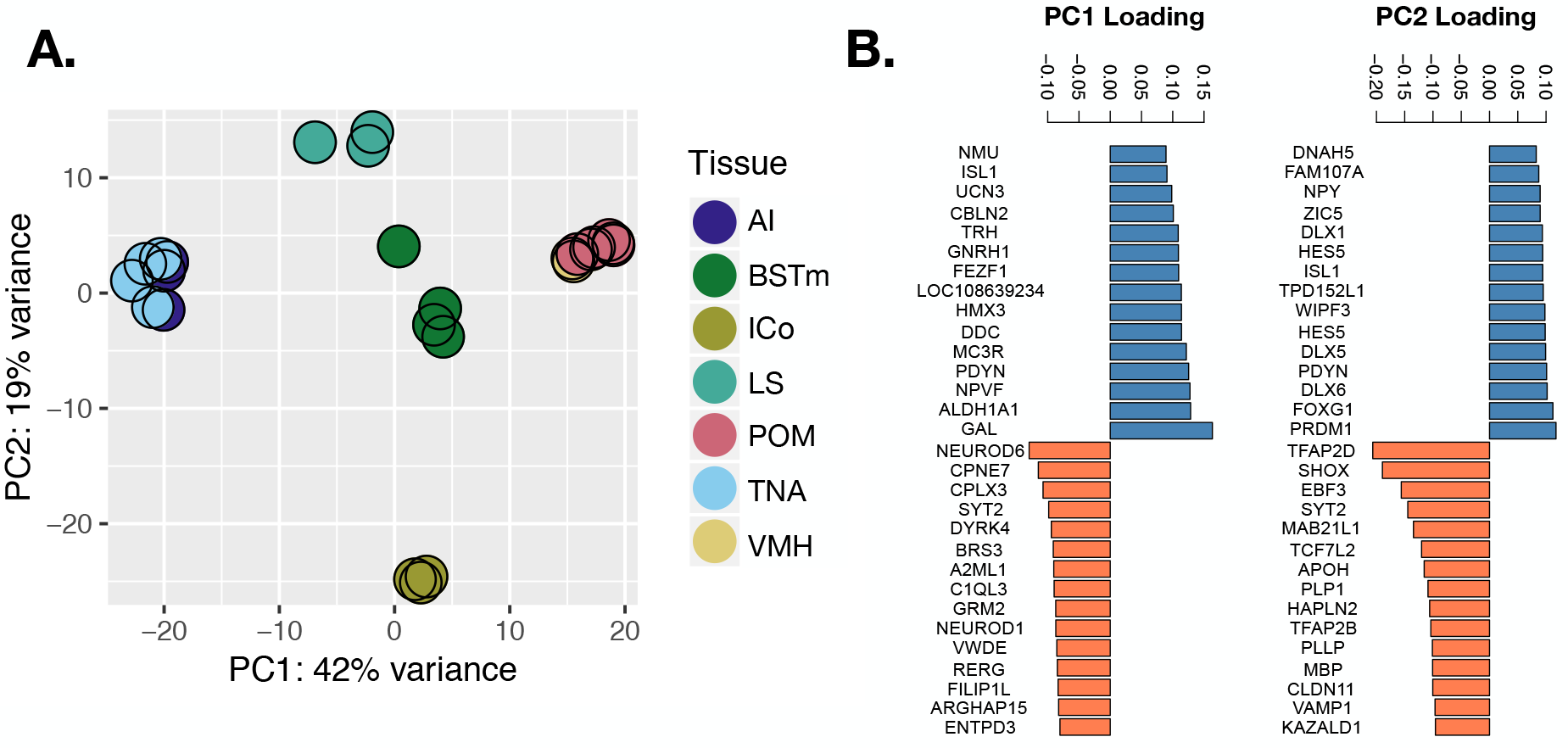
**A)** Scatterplot of Principal Component (PC) 1 versus PC2 and **B)** top 30 genes b with based on loadings on each PC.

### Associations Between Expression Networks and Brain Nuclei

Using the analysis parameters described above, WGNCA analysis constructed a total of 19 modules (Fig. S1-S2). Sixteen modules showed expression patterns that were associated with at least one brain region in the SBN (see Fig. S1). Based on our chosen module dissimilarity threshold, we retained some modules that showed similar expression patterns. For example, genes in the blue, black and white modules have relatively high expression in amygdala (TnA, Ai) and low expression in hypothalamus (VMH, POM), whereas genes in the brown, dark grey and light cyan modules have the opposite expression pattern (Fig. 2). Although more stringent merging of modules would have joined some of these modules, we retained these modules as distinct because they also reflect subtle and potentially important differences in regional patterns of gene co-expression (Fig. 2). These differences were particularly apparent for genes with high module membership in their assigned module. For example, genes with high module membership in the black module (e.g., progesterone receptor (PGR)) are distinguished from those assigned to the blue module (e.g., proenkephalin (PENK)) by patterns of high and low expression in LS, respectively (Figure 2). Likewise, genes with high module membership in the brown module are differentiated from those assigned to the dark grey module by having low expression in LS. Module assignment for genes with lower module memberships, however, can be considered more ambiguous (despite statistical assignment to a particular module). An example of this is the gene for estrogen receptor alpha (ESR1), which was assigned to the blue module even though it has a higher estimated module membership for the black module (MMblue = 0.68 vs MMblack = 0.80, see Figure 3). Such patterns reflect the lack of orthogonality in WGCNA modules; that is, a given gene can be associated with multiple regulatory networks. Three modules, including the “grey” module which comprises the set of unclustered genes (20 genes) showed significant variation among individual birds (mediumpurple3, *r* = 0.64, *p* = 6e−04; darkolivegreen, r = 0.81, *p* = 9e−07).

**Figure 2.**
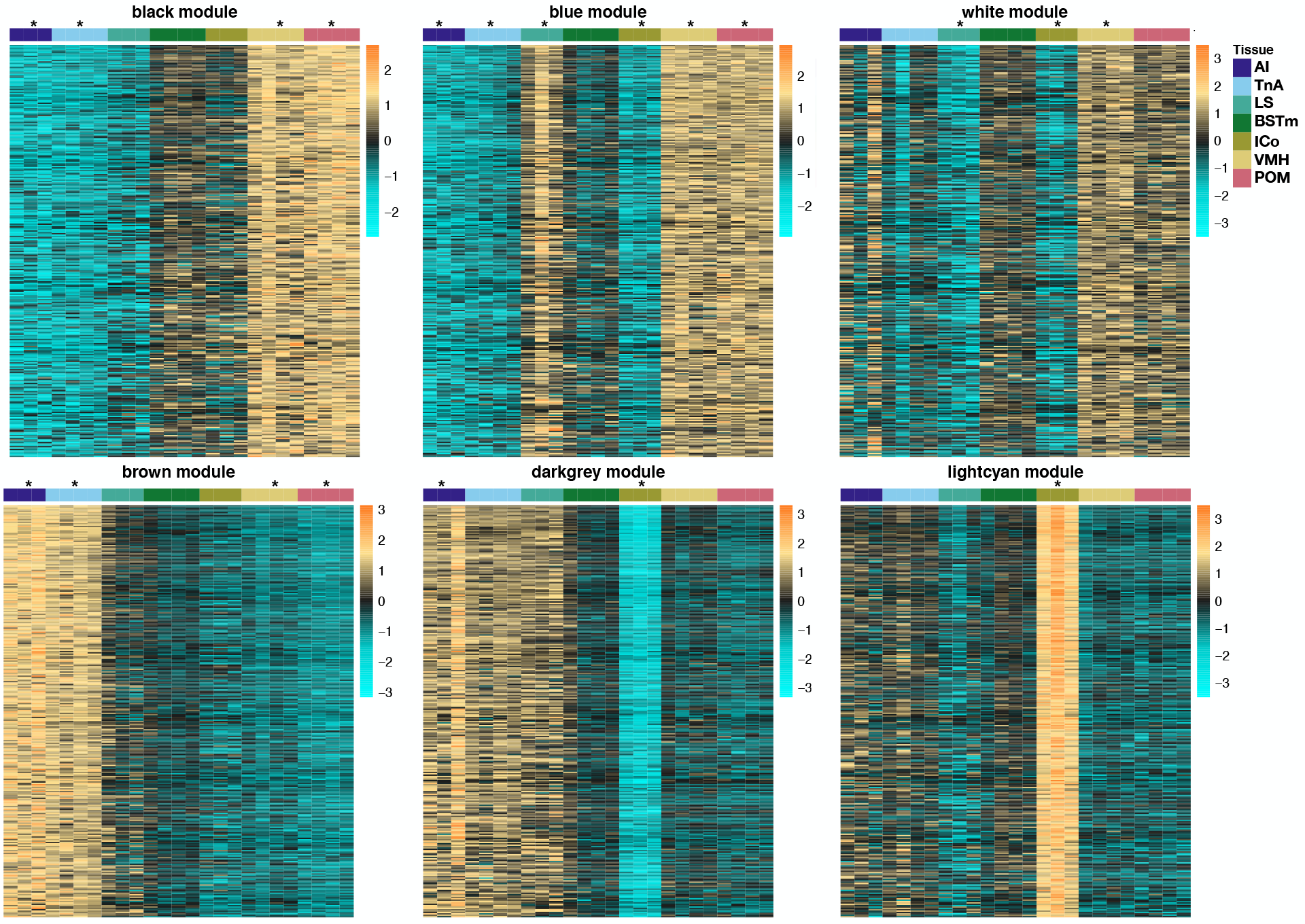
Heatmaps of top 300 genes associated with each of six modules (out of 19 total). Modules were selected for display based on their association with genes of interest (Table 1, Figure 3) and to highlight regional differences in gene expression. Asterisks indicate statistical associations (*p* < 0.05) between a module and brain region (see also Figure S1).

### Examination of Neuroendocrine Candidate Genes

To highlight the relationship between WGCNA modules and genes related to neuroendocrine signaling and behavior, we compiled a list of genes encoding key hormones, steroidogenic enzymes, and hormone receptors (Table 1, Fig. 3). In doing so we observed that the majority of the well-studied neuropeptides and hormone receptors (e.g., VIP, DRD1, ESR1, AR) were assigned to just four modules, blue, black, brown and dark grey, representing the two opposing gradients of expression described above. For example, androgen receptor (AR), estrogen receptor alpha (ESR1), progesterone receptor (PGR) and vasoactive intestinal polypeptide (VIP) show high expression in the hypothalamic nuclei (POM and VMH) and low expression in the amygdala in both the blue and black modules (Table 1, Fig. 2–3). By contrast, the brown and dark-grey modules had low expression for these same genes in hypothalamic nuclei but high expression in the amygdala (Figure 2). These four modules also included a number of serotonin and dopamine receptors, as well as insulin receptor and the cannabinoid receptor CNR1 (Table 1).

Other candidate neuroendocrine genes were distributed among five other co-expression modules (Table 1). In general, these genes had lower module membership values (MM = 0.32 − 0.77), indicating lower specificity of expression among sampled brain regions. Among these are two genes involved in steroid metabolism, including 5-alpha reductase (SRD25A) and aromatase assigned to the light cyan (MM = 0.41) and white (MM = 0.77) modules respectively. Two key nonapeptide hormone-encoding genes, OXT (encoding oxytocin and neurophysin1) and AVP (encoding arginine vasopressin, neurophysin 2 and copeptin) were not annotated in the *M. vitenillus* genome, and thus we do not have information on their expression. We focus the remainder of our analysis on the six modules within which candidate genes were most strongly represented (Fig. 2).

Analysis of regional variation in gene expression across SBN regions also enables us to describe patterns among paralogous hormone receptors. Gene duplication and subsequent sub- or neofunctionalization provides a mechanism by which hormonal action can be modulated to have context-specific and region-specific effects^61,62^, and thus has played an important role in the evolution of hormonal signaling^63,64^. We specifically examined paralogous receptor genes for estrogen, galanin, vasoactive intestinal peptide, serotonin, neuropeptide Y, dopamine, somatostatin and GABA (Table 1). All of these gene families except for the GABA receptors (R1, R3, B3) included members that that were assigned to different modules indicating some degree of expression divergence for each gene family.

**Figure 3.**
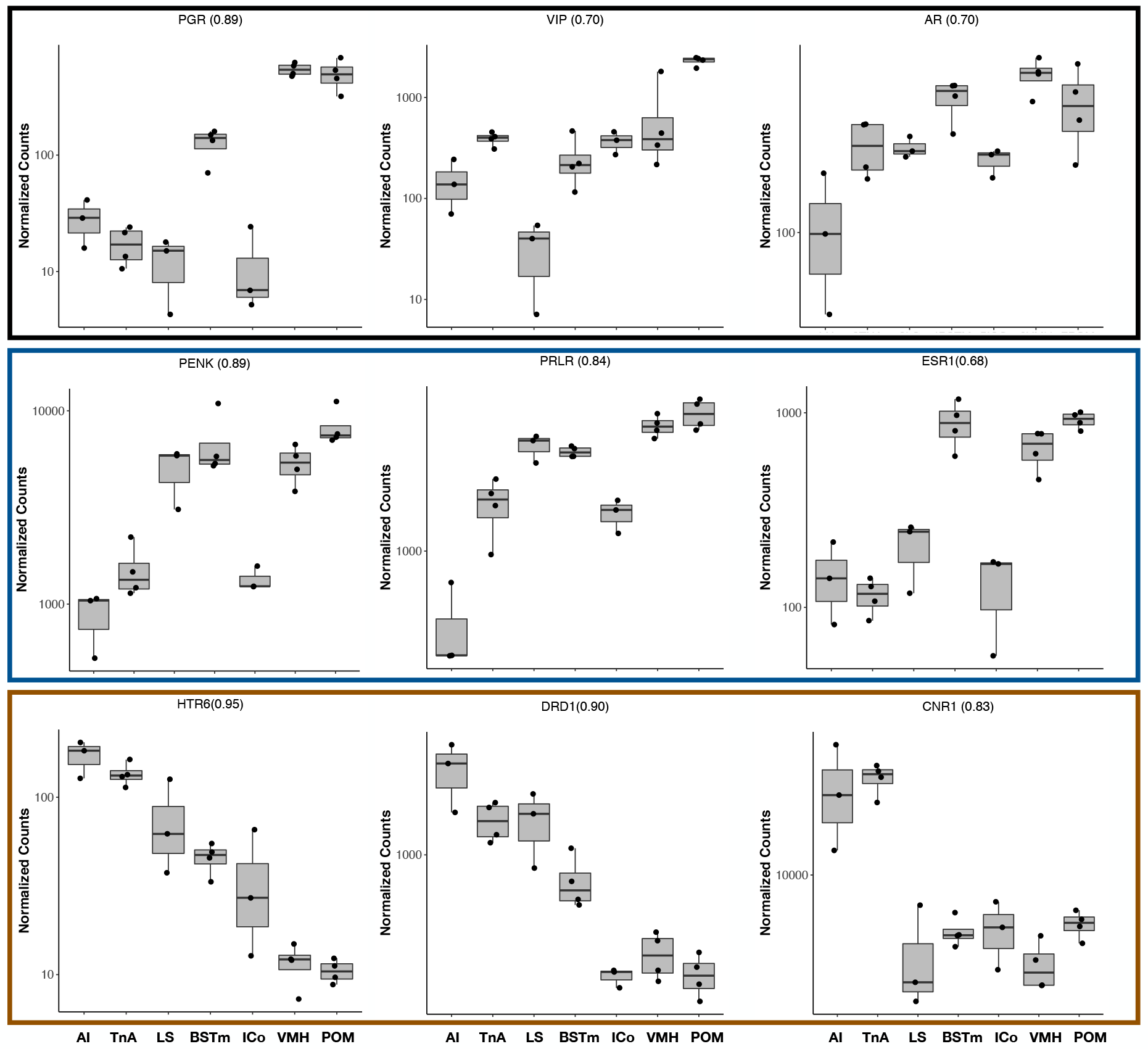
Expression plots (normalized read counts) for candidate genes associated with three key modules, blue, black and brown. Module assignment is denoted by the colored box around each panel of three genes. Module membership for each gene is presented within parentheses besides each gene name.

### Functional Annotation of WGCNA Modules

In addition to the candidate gene approach, we used Gene Ontology analysis to systematically characterize WGCNA modules in terms of function. These analyses revealed two broad themes in terms of functional enrichment. As might be expect based on the candidate gene analysis, we observe enrichment for GO categories related to neuropeptide signaling and hormone receptor activity across multiple modules (Tables S1-6). As in the PCA analysis presented above, we also observed an enrichment of terms related to developmental processes and tissue differentiation in the brain.

As the modules themselves have correlated expression patterns, GO analyses based on module memberships were similar. The white and black modules (high hypothalamic expression but low expression in the amygdala) showed significant enrichment for multiple GO categories related to neuropeptide signaling (e.g., “neuropeptide signaling pathway”, white: *p* = 3.22e−04, black: *p* = 0.04; “dopamine metabolic process”, white, *p* = 0.04, Tables S1, S2). The black module also showed an enrichment for genes involved in “developmental process” (*p* = 0.02) including the orthopedia homeobox gene (OTP, MM = 0.91). This module was also enriched for genes in the “smoothened pathway” (*p* = 0.02), including the classic developmental morphogen sonic hedgehog (SHH, MM = 0.80) and its receptors patched 1 (PTCH1, MM = 0.83) and PTCH2 (MM = 0.82). Although some of our candidate neuroendocrine-related genes mapped to the blue module (Table 1), this module was primarily enriched for GO categories related to development and regional differentiation (e.g., “pattern specification” (*p* = 3.0E−11) and “determination left/right symmetry” (*p* = 6.5E−09), Table S2). The strongest enrichments in the blue module were a large set of categories related to cilium morphology, which may relate to aspects of cellular structure and compositional differences among brain regions.

Among the modules that had high expression in the amygdala (TnA) and AI, both the brown and dark grey modules (Tables S3, S4) were significantly enriched for GO categories relating to synaptic transmission, plasticity (dark grey, *p* = 3.5e-06), and learning and memory (dark grey, *p* = 7.8e−04). These categories include dopamine receptors DRD1 and DRD5, cannabinoid receptor CNR1, and a suite a glutamate receptors (GRIN2A, GRIN2B, GRM2, GRIA2 and GRIA3). The dark grey module also included signatures of development-related genes, including those associated with “positive regulation of dendritic spine development” (*p* = 3.14e−04). Although the light cyan and dark grey modules had similar expression profiles in the hypothalamus and amygdala, genes expression patterns in the LS differed markedly between these modules, with low expression for light cyan genes but high expression for dark grey genes (Fig. 2). Genes strongly associated with the light cyan module (Table S5) were highly enriched for functions related to cellular energetics (e.g., “mitochondrial electron transport, NADH to ubiquinone” (*p* = 5.4e−12), “generation of precursor metabolites and energy”, *p* = 6.5e−12).

Two modules with the strongest enrichment for behavior-related genes were the black and dark grey modules. Thus, we further examined the topology of these networks to better characterize potential regulatory interactions. In the black module, the most highly connected hub gene encodes the synaptic organizer protein CBLN2 (98 connections, Fig. 4). Two other key hubs were serine/threonine kinase 32B (STK32B, 68 connections) and Dopa decarboxylase (DDC, 21 connections; Fig. 4). DDC also had the highest module membership among genes in the black module. In the dark grey module, the gene with highest connectivity was Signal Induced Proliferation Associated 1 like 1(SIPA1L1, 95 connections among the top 500 genes). Additional highly connected genes in this module include a series of glutamate-related genes (GRIN2B, SHANK2, DLGAP1& DLGAP2).

**Figure 4.**
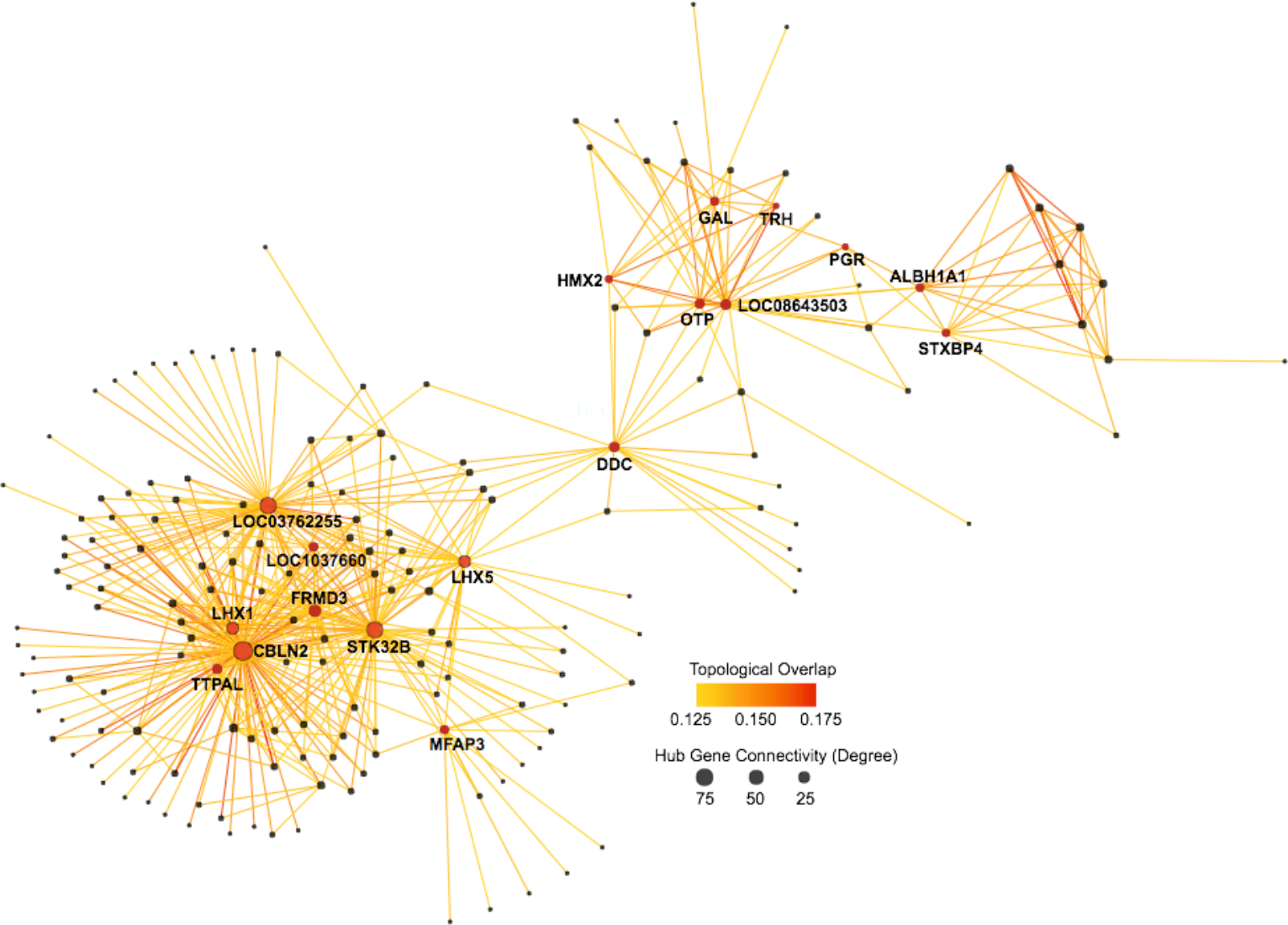
Network diagram of top 500 genes in the black module. Gene names are shown only for hub genes (>20 connections).

## DISCUSSION

We have provided here a fundamental picture of heterogeneity in gene expression across several nodes of the vertebrate SBN. This heterogeneity reveals composite signatures of gene co-expression that reflect differences in cellular composition and activity among brain regions. These signatures highlight regional differences in these mechanisms across nodes of the SBN. Overall, our results reveal opposing gradients of expression among gene modules, whereby some suites of genes are highly expressed in the hypothalamic nuclei and lowly expressed in the amygdala, or vice versa. Genes tightly associated with 19 defined regulatory modules reveal distinctive expression signatures that differentiate each of the sampled SBN nodes and adjacent brain regions.

The majority of candidate loci encoding neuropeptide hormones, steroid and neuropeptide hormone receptors, and steroidogenic enzymes were associated with modules that showed high hypothalamic expression and low amygdalar expression (i.e., black, blue, and white modules; Fig. 2). This finding underscores that high levels of neuropeptide activity and steroid sensitivity in some SBN brain regions play integral roles in how neuropeptide and steroid signaling modulate vertebrate social behavior (e.g., Goodson 2005). A subset of serotonin, dopamine and cannabinoid receptors associated with the brown module show the opposite pattern, with high expression in the amygdala (TnA) and an adjacent nucleus (AI) and low expression in hypothalamic nuclei. This pattern likely reflects the role of these genes and brain regions in reward circuitry.^65^ The amygdala nucleus (TnA) and AI were also enriched for genes involved in synaptic plasticity, especially for genes associated with glutamate receptor activity (e.g., GRIN2A, GRIN2B, FMR1, TRPM1, HOMER1, GRIA3, GRIA2, GRIN3A). Across the SBN, we observed striking variation in the distribution of various transmembrane receptors (e.g., ion-channel genes), a pattern similar to that observed in an analysis of variation in gene across the song control system in songbirds.^31^

Dopa decarboxylase (DDC) was the gene with the highest module membership in the black module, where it was a highly interconnected hub gene (21 of the 500 strongest connections, Fig. 4). DDC is an enzyme that catalyzes the production of dopamine and serotonin from their respective precursors, tyrosine and tryptophan. Indeed, the constructed network connected DDC with GTP cyclohydrolase 1 (GCH1), another enzyme involved in the production of dopamine and serotonin. As a module hub gene, DDC also linked the “neuro-endocrine” and “developmental” components of this regulatory network. On the neuroendocrine side, DDC has direct links to genes for progesterone receptor (PGR), thyrotropin-releasing hormone receptor (TRHR), galanin and galanin message-associated peptide (GAL), and a ghrelin receptor (GHSR). DDC, however, is also linked to genes with developmental functions like LIM Homeobox 5 (LHX5, also a hub, 29/500 strongest connections), SHH, and OTP among others. The most highly interconnected gene within the black module was cerebellin2 (CBLN2), a gene with known roles in the structural organization of synapses.^66^ SSH as well as other LIM homeobox genes (LHX6/7) have previously been described as markers of nodes within the mesolimbic reward system and/or SBN.^6^

The neuropeptide gene with the highest membership in any module was galanin (GAL, 14 connections, MMblack = 0.91; Fig.4). Galanin and its receptors are known to be abundant in the hypothalamus and preoptic area of the mammal brain^67^, whereby galanin actions may influence dopamine, oxytocin, and gonadotropin-releasing hormone (GnRH) release.^68^ Galanin is also believed to play a role in the mesolimbic reward system.^69^ Consistent with its known pleiotropy, GAL, like DDC, holds a key node in the black regulatory module (14 connections, MM = 0.98). GAL is directly linked to PGR, thyrotropin-releasing hormone (TRH), TRHR, GHSR, SSH and OTP in the black module. These connections between GAL, DDC and OTP are especially interesting because of the known role of OTP in neuronal differentiation. Both OTP and Nescient Helix-Loop-Helix 2 (NHLH2, also assigned to the black module, MM = 0.91) play a role in the specification of POMC (Pro-opiomelanocortin), NPY, AGRP (Agouti Related Neuropeptide), GnRH, and dopaminergic neurons within the hypothalamus.^70,71^ Likewise OTP and POU3F2 (POU Class 3 Homeobox 2, not annotated in our dataset) are required for the expression of oxytocin, vasopressin, TRH and corticotropin-releasing hormone (CRH) in hypothalamic neurons.^70^ Thus, our reconstructed network supports known regulatory interactions between OTP and multiple neurohormones.

Most of the examined neuropeptide receptor gene families examined include paralogs that were assigned to different modules. This finding supports an evolutionary model of gene regulatory divergence following gene duplication. One exception to this rule was that all three detected GABA receptors were assigned to the same module (Table 1). Likewise, ESR1 and GALR3 were assigned to the black module, whereas ESR2 and GALR1 were assigned to the blue module. As mentioned previously, black and blue modules show parallel expression patterns so paralogous estrogen and galanin receptors do as well. By contrast, more striking differences in expression (i.e. assignment to modules with contrasting expression patterns) is observed for VIPR1 versus VIPR2 and NPY1R versus NPY2R and NPY3R. A particularly interesting case are the DRD1 and DRD2 dopamine receptors, which are assigned to brown and blue modules, respectively. DRD1 and DRD2 have independently evolved the ability to bind dopamine from serotonin receptors^72^. DRD1 is more closely related to the serotonin receptors 5HT4 and 5HT6, and shares a more similar expression pattern with these serotonin receptors than it does DRD2. DRD2, on the other hand, shares a more similar expression pattern to serotonin receptor 5HT2, to which it is more closely related. Comparative studies across species with varying gene complements could reveal the history of possible neo- and sub- functionalization of these genes.

Viewed cumulatively, our analyses contribute new evidence to a growing body of research revealing the heterogeneity of gene expression in the brain. By linking this expression heterogeneity to the functionally integrated nuclei within the vertebrate SBN, we hope that our results will facilitate subsequent neuroendocrine and genomic studies of the molecular mechanisms regulating social behavior. Although our current sample size prohibits a useful comparison among male social classes in these manakins, our study lays the foundation for future work on how variation in neural gene expression across nodes of the SBN may underlie individual differences in behavioral phenotype and, ultimately, male social status. Importantly, our work highlights the feasibility of detailed neurogenomic analysis in a non-model system studied in a field setting, and iterates the importance of examining neuroendocrine gene expression across multiple nodes of the SBN to ultimately understand links between genes and behavior. Such comprehensive analyses that include resolution at the nucleus level should improve the interpretability of neurogenomic studies going forward, and encourage analyses of diverse systems in natural settings that can yield a broader understanding of the mechanisms underlying vertebrate social behavior. Finally, these results suggest that differences in the degree to which co-expressed neuroendocrine genes are up and down regulated across nodes of SBN may underlie among-individual differences in social behavior.

**Table 1.**
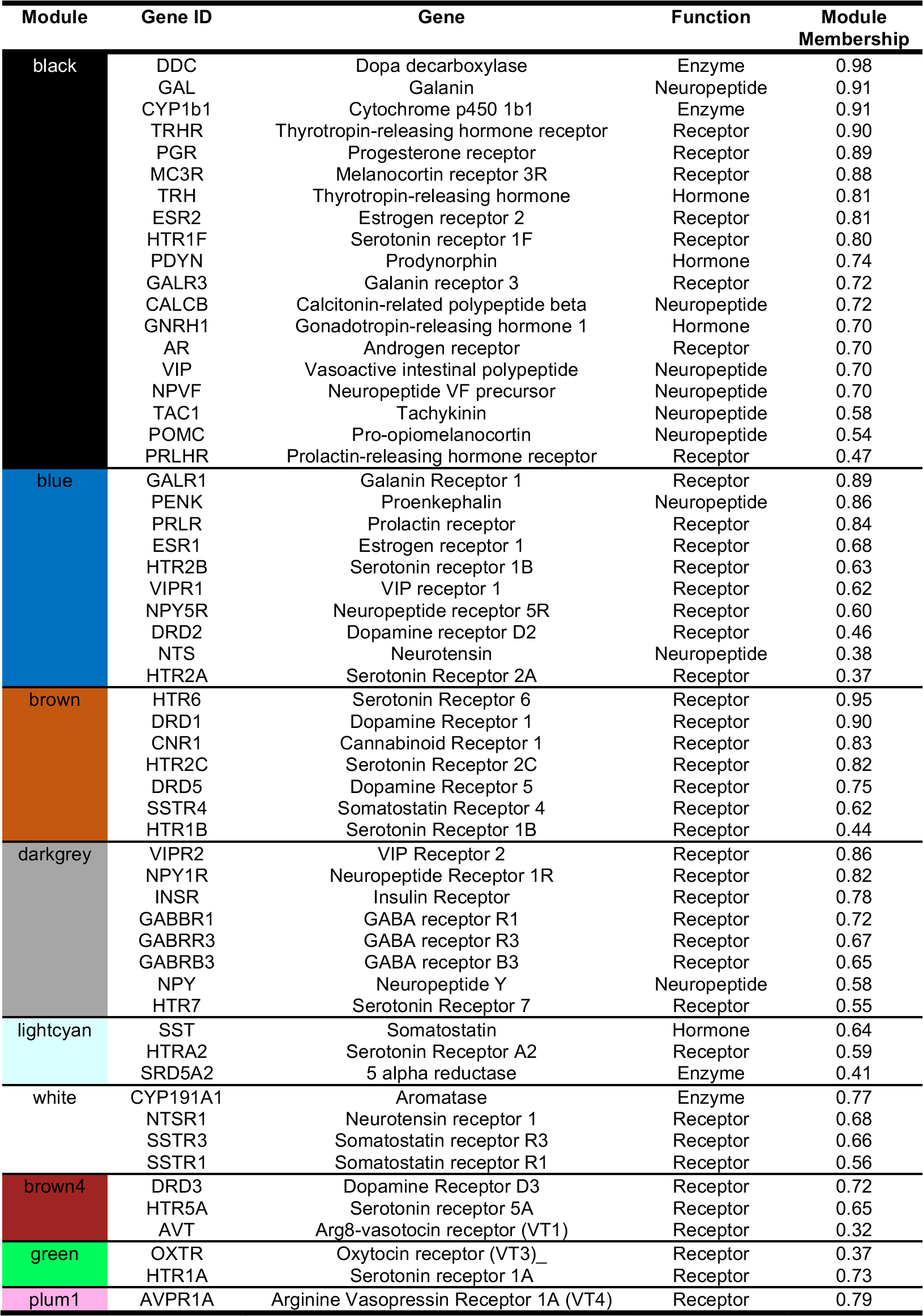
Module assignment for key hormones, receptors and enzymes.

## AUTHOR CONTRIBUTIONS

**BMH** designed the study, conducted field and lab work and wrote the manuscript

**TBR** designed the study, conducted field work, assisted with analyses and wrote the manuscript

**ITM** designed the study, conducted field work, and provided feedback on the manuscript

**CNB** designed the study, analyzed the RNA-seq data and wrote the manuscript

## ACKNOWLEDGEMENTS

Thanks to David Clayton and Daniel Newhouse for providing feedback on drafts of this manuscript. Daniel Newhouse also provided assistance with code for WGCNA analyses. Matthew Fuxjager and Susan Fahrbach provided access to lab space and equipment at Wake Forest University, where cryosectioning, microdissection, and RNA extraction were carried out. Collection of birds was done in accordance with Smithsonian ACUC #14-25 & 17-11, export and import permits included MAE-DNB-CM-2015-0008, 006-016-EXP-1C-FAU-DNB/MA & USDA APHIS 126133. The development of this work was facilitated by a NSF Research Coordination Network grant (1457541). Funding for this work was provided by NSF 1353085 (TBR, ITM & BMH) & 1456612 (CNB).

---

**Figure S1.**
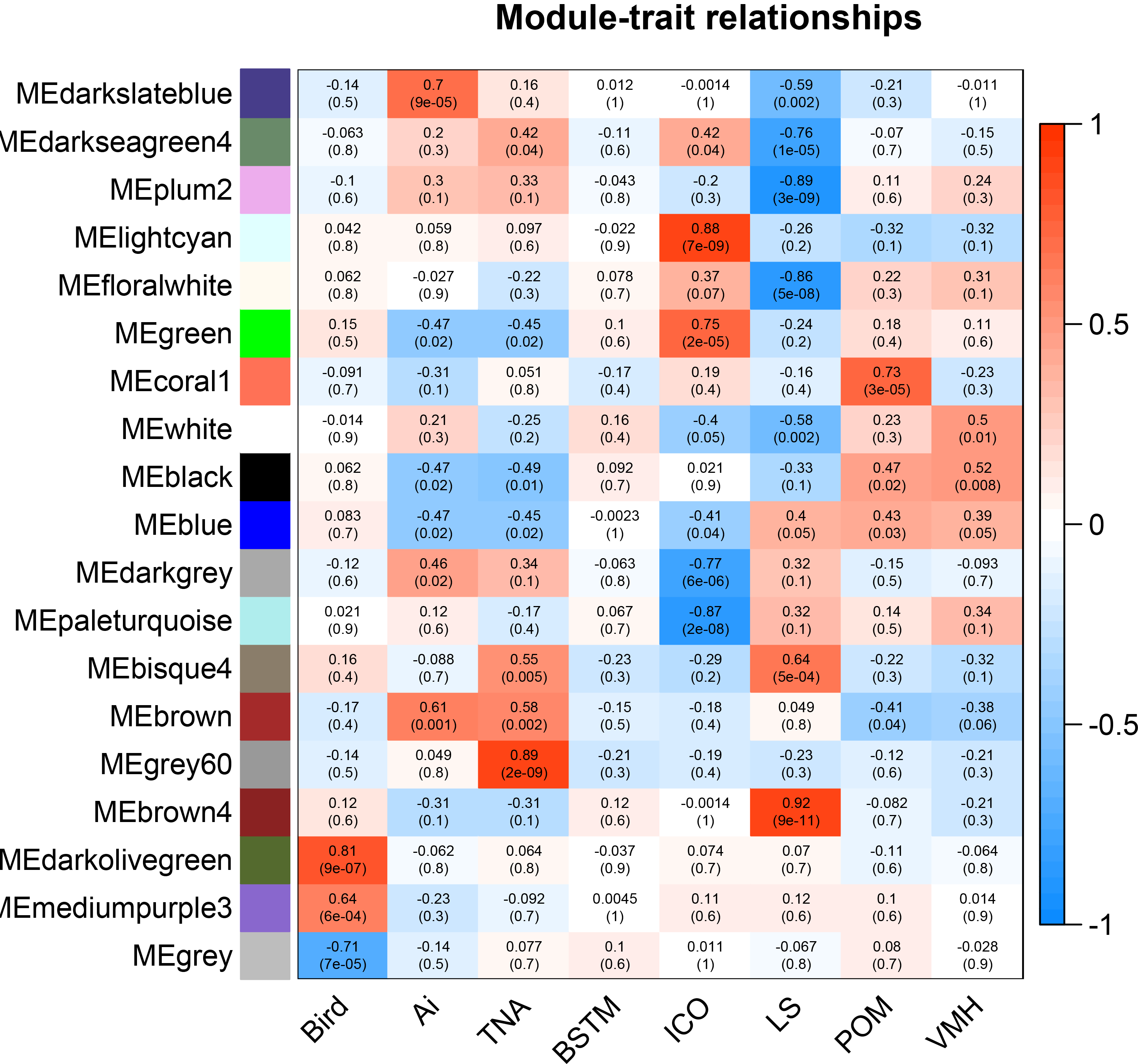

**Figure S2.**
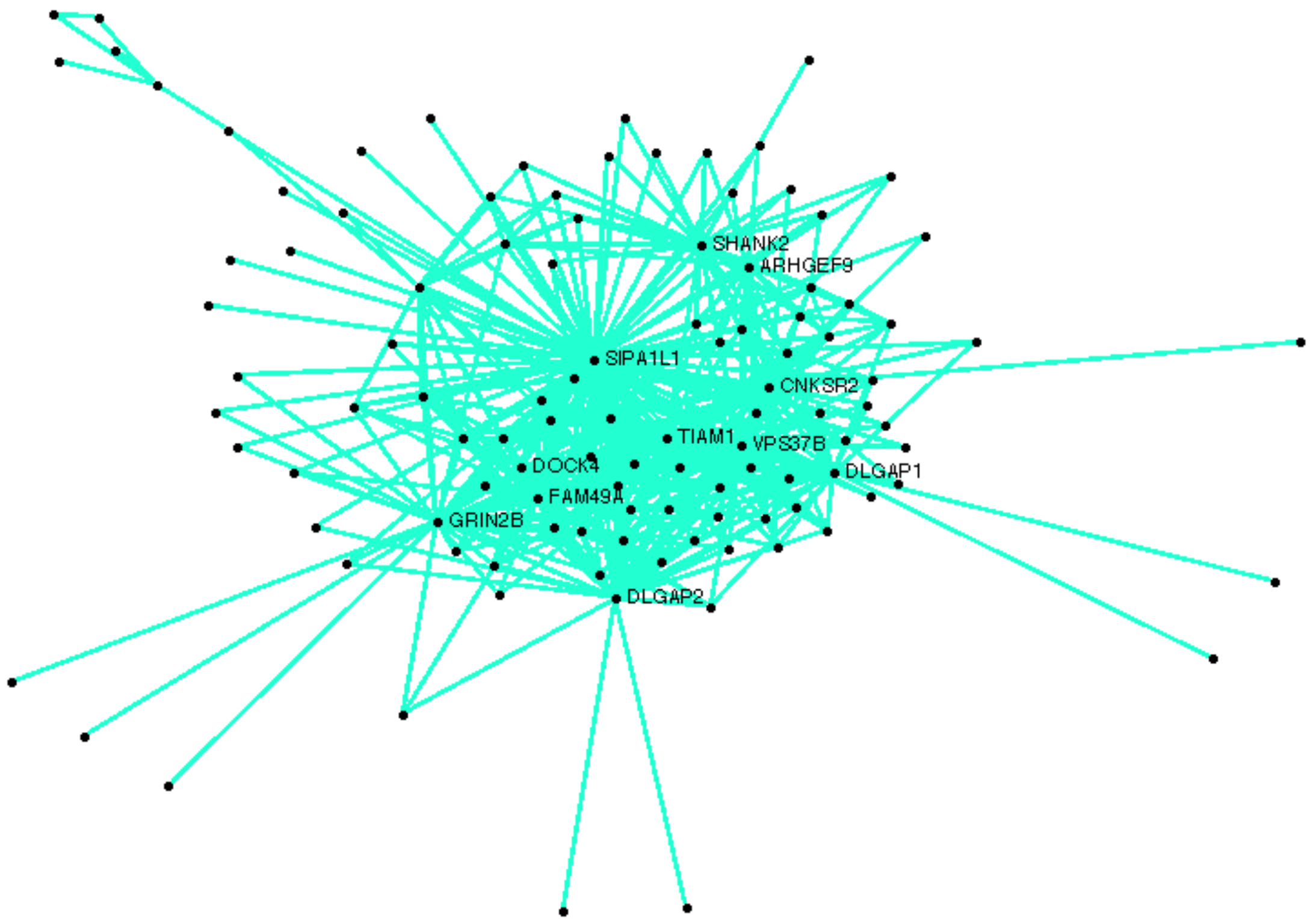

